## Supplemental files for "Single nucleotide polymorphism induces divergent dynamic patterns in CYP3A5: a microsecond scale biomolecular simulation of variants identified in Sub-Saharan African populations"

#### 1 Assessing the convergence of the molecular dynamics trajectory

To assess the convergence of each trajectory we have made a clustering analysis over all the snapshots using cpptraj. We choose to make a hierarchical clustering using a cutoff value of 2 Å. The cumulative number of clusters as a function of time was calculated as well as the evolution of informational entropy according to the following formula.

$$H = -\sum_{i=1}^{n} p_i log(p_i)$$

where $p_i$ is the probability of the $i^{th}$ found cluster, as a function of simulation time.

The wild type form, Y53C, I149T, and I276T had shown a constant number

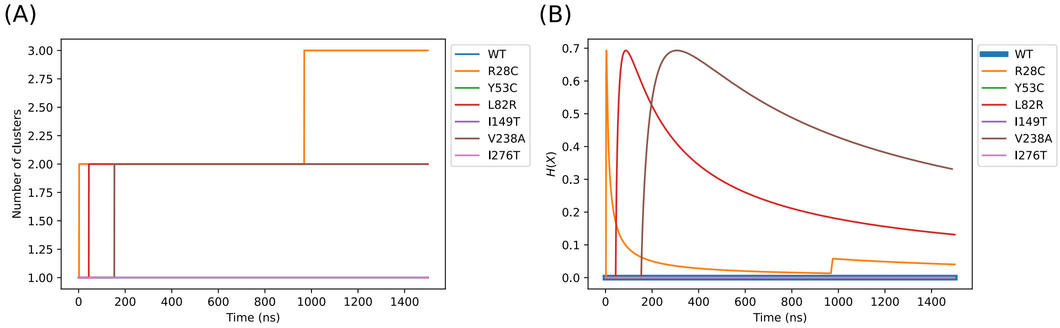


Figure 1: Analysis of the molecular dynamics simulation using the cumulated number of clusters (A) and the onstancy of the cluster entropy

of clusters of 1 since the beginning of the simulation as well as an entropy value of 0. R28C, L82R, and V238A have not converged properly indicating a likely shift in the conformational space of these variants compared to the wild-type form. Since we are comparing the effect to the wild-type form, this is acceptable. However, we notice from the entropy calculation that the trajectories tend to converge at the end of the simulation.

#### 2 Fluactuation of CYP3A5 residues

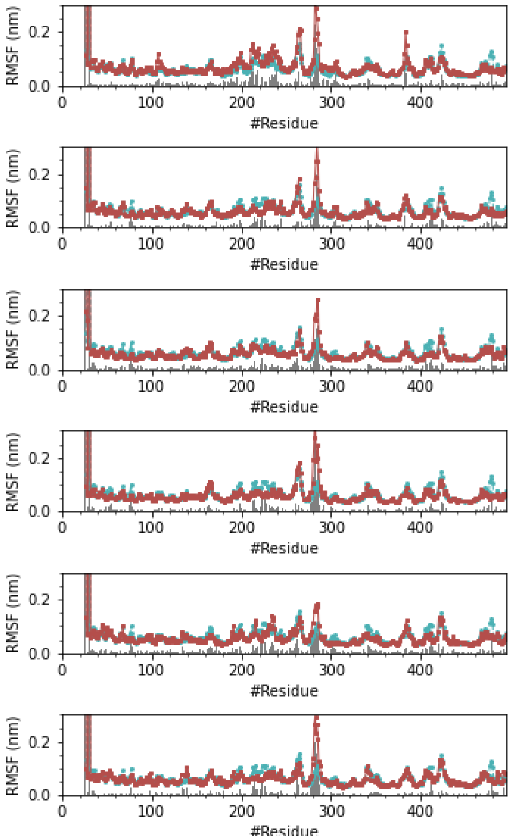


Figure 2: Root Mean Square Fluctuation of CYP3A5 residues. From top to bottom: WT, R28C, Y53C, L82R, I149T, V238A and I276T.

# 3 PC3 Vs PC4

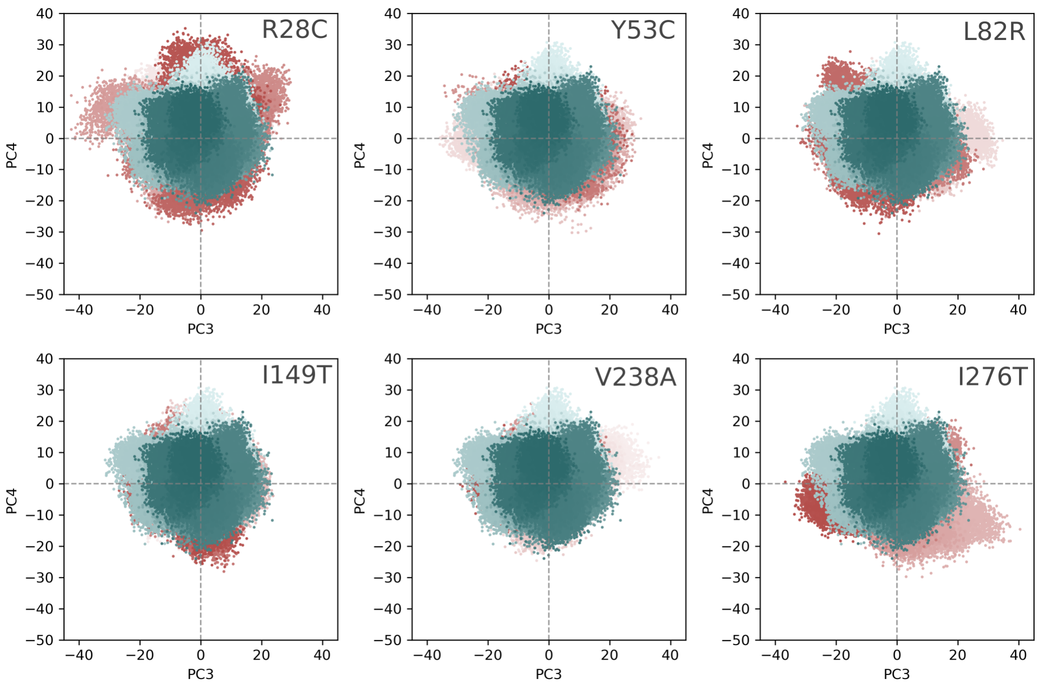


Figure 3: Principle component analysis, PC3 Vs PC4

### 4 Free energy landscape of all CYP3A5 variants

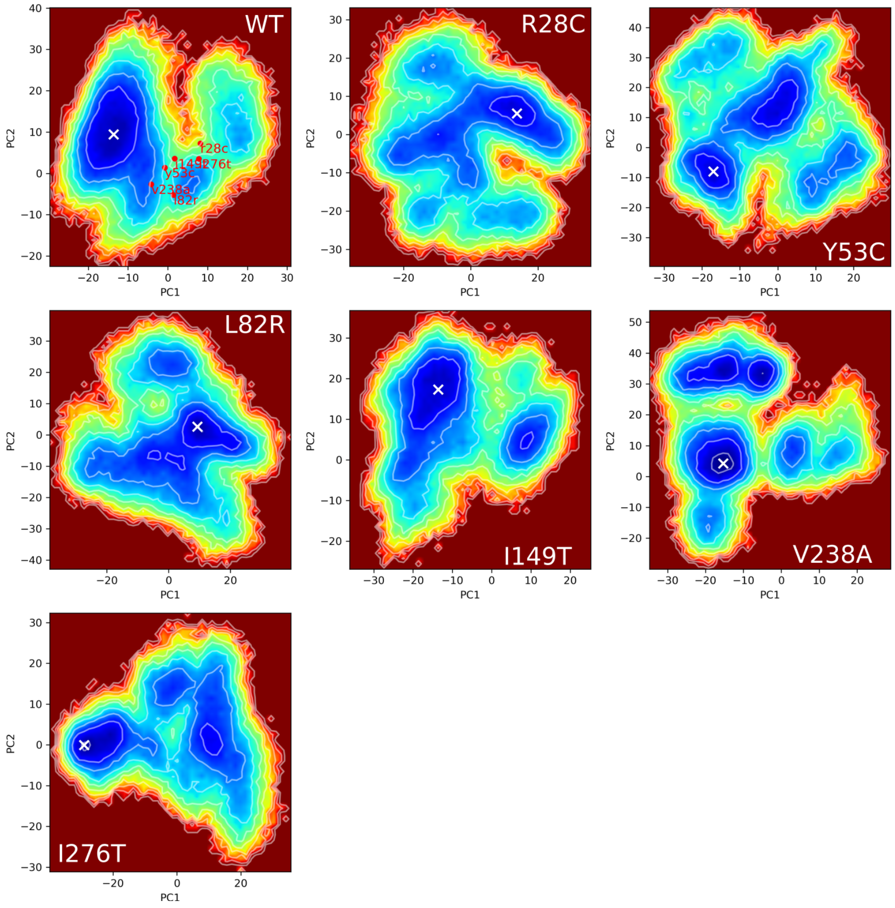


Figure 4: Free energy landscape of the wild type and CYP3A5 variants characterized from sub-Saharan African population. The white X marker indicates the position of the global minima. The position of the conformations at the local minimum of each variant was projected onto the free energy landscape of the wild-type form.

### 5 Convergence of MM-GBSA

The convergence of the MM-GBSA calculation was verified for the complex of CYP3A5/ritonavir for the wild type form for a total simulation time of 300 ns. We plotted the MM-GBSA energy over an increasing time step of 5 ns in two fashions: forward (all the frames below the upper boundary of the intervals are considered in the calculation) and backward (all the frames above the lower boundary of the interval are considered in the calculation). We can see from the plot that the energy converges at 100 ns simulation time both for the forward and backward calculations.

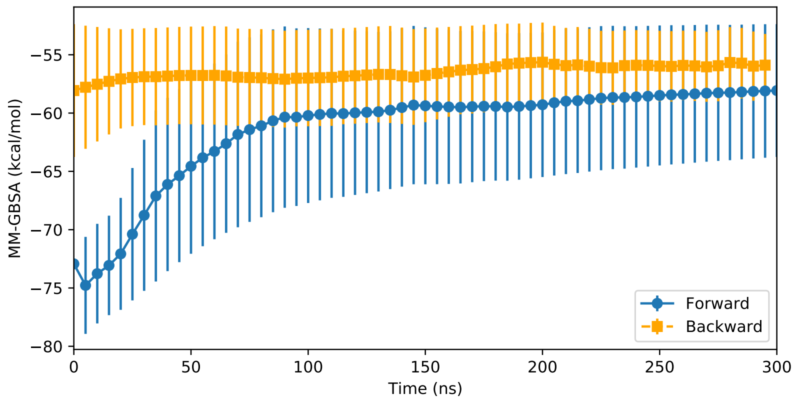


Figure 5: Convergence of the MM-GBSA energy for a simulation time of 300 ns calculated for the complex CYP3A5/ritonavir for the wild type form.
